## Supplementary Information for "OpenCap: 3D human movement dynamics from smartphone videos"

### Methods

##### Intra- and inter-device analysis of camera intrinsic parameters

OpenCap includes a database that maps iOS device models to camera intrinsic parameters (principal point, focal length, and distortion parameters). The intrinsic parameters were computed for each model with a precision-manufactured 720x540 mm checkerboard with nine rows, 12 columns, and 60 mm squares.

To evaluate the reliability of the algorithm for estimating the intrinsic parameters (i.e., intra-device testing), we computed the intrinsic parameters five times using the same device (iPhone 12 Pro), but each time based on a different set of 50 images of the checkerboard. The standard deviation across the five calibrations, expressed in percent of the mean, for both the focal length and the principal point was lower than 1%, suggesting a low sensitivity of the intrinsic parameters to the images used for their estimation and thus a high reliability of our algorithm.

To evaluate whether intrinsic parameters are sensitive to the device used for their estimation (i.e., inter-device testing), we computed the intrinsic parameters with five devices of the same model (iPhone 12 Pro). The standard deviation across the five devices, expressed in percent of the mean, for both the focal length and the principal point was lower than 1%, suggesting that the camera hardware across devices of the same model is consistent and thus that intrinsic parameters computed with one device can be used for another device of the same model.

##### Comparison of kinematics using a small printed versus a large, high-precision checkerboard

To determine whether a checkerboard printed with a standard printer is sufficient for calibrating camera extrinsic parameters in OpenCap, we compared marker positions and kinematics computed when the cameras are calibrated with the printed checkerboard to when they are calibrated with a large precision-manufactured board. The printed checkerboard was printed on A4 paper (210x175 mm, five rows, six columns, 35 mm square size) and taped to plexiglass, and the precision-manufactured checkerboard (Calib.io, *Svendborg*, Denmark) was 720x540 mm (nine rows, 12 columns, and 60 mm squares).

With two cameras placed at 45°, we recorded a calibration video of both checkerboards. One individual then performed a standing neutral trial followed by a squat trial (five repetitions). We calibrated the cameras using both checkerboards, then computed the difference in marker positions and kinematics computed with each calibration using the OpenCap pipeline (Methods: Design). The mean per marker difference for the 20 3D video keypoints was 3 mm and the mean absolute error of kinematics was less than 1° and less than 1 mm for rotational and translational degrees of freedom, respectively. These results demonstrate that calibrating with a checkerboard printed with a standard printer does not impact marker position and kinematic results, and that a printed board can be used with near-identical accuracy as the precision-manufactured board used during the in-laboratory validation portion of our study.

##### Video collection and pose estimation: 2D keypoint pre-processing

OpenCap includes several pre-processing steps to improve the fidelity of the 2D video keypoints prior to triangulation. First, many pose detection algorithms, like OpenPose, do not track the same person between frames. We implemented a person tracker using bounding boxes around the keypoints of each person identified by OpenPose. The largest bounding box of any video frame is identified as the person of interest, and this box is tracked between frames until the between-frame change in bounding box corner positions exceeds an empirically determined threshold. Second, occlusion of leg markers is common during activities like gait. OpenCap identifies occlusions using the relative confidence scores of the right and left leg markers and uses cubic splines to replace the occluded keypoint positions. Finally, to reduce high-frequency noise in the keypoint positions between frames, OpenCap filters keypoint positions using fourth-order, zero-lag Butterworth filters (12 Hz for gait trials, 30 Hz for non-gait trials; see Video collection and pose estimation: Synchronization below).

**Video collection and pose estimation: Synchronization**

The connection between the iOS devices and the web application is internet based and video recording is therefore not precisely synchronized. We developed two custom algorithms to synchronize recorded videos using keypoint trajectories: one for gait and another for all other activities. OpenCap identifies gait trials by cross correlating the speeds (in the image plane) of the right and left ankle keypoints; if the signals have a large maximum cross correlation and a time delay of 0.1–1 s, the trial is deemed a gait trial. The time delay for gait trials is computed using the sum of cross correlations between the right and left ankle and heel marker speeds between two videos. OpenCap triangulates the keypoints at the three time delays closest to zero, and selects the delay that corresponds to the lowest error between reprojected 3D keypoints and 2D video keypoints. For non-gait trials, OpenCap selects the time delay corresponding to the maximum cross correlation of the summed vertical speed (in the image plane) of all keypoints between cameras.

**3D marker set augmentation**

We trained two LSTM networks: an *arm model* to predict the positions of eight arm markers from the positions of nine arm and torso keypoints, and a *body model* to predict the positions of 35 body markers from the positions of 13 lower-limb and torso keypoints. Both models also include height and weight as inputs. To generate a training set for these networks, we synthesized corresponding pairs of 3D video keypoints and 3D anatomical markers from 108 hours of motion capture data processed in OpenSim[1]. Note that since not all datasets included arm data, we only included 79 hours of motion capture data (68 subjects from 5 datasets) to train the *arm model*.

To build the training set of synthetic data, we first combined 10 existing datasets (336 subjects) containing scaled OpenSim models and motion data (i.e., results from inverse kinematics) from published biomechanics studies[2–11]. For each dataset, we split the data in a training set (~80%), validation set (~10%), and test set (~10%). We performed the splitting on a per-subject basis, such that data from a subject was not part of multiple sets. We then added virtual markers to the scaled OpenSim models corresponding to the video keypoints and the anatomical markers that we aimed to predict with our LSTM networks. We positioned the video keypoints at the joint centers. We positioned the anatomical markers based on a standard marker-based motion capture protocol[12]. Next, for each time frame of each motion file, we extracted the 3D positions of each virtual marker using OpenSim Point Kinematics tool. To augment the dataset to include shorter and taller subjects, we repeated this process after uniformly scaling each OpenSim model by 90, 95, 105, and 110%. We then expressed the 3D positions of each marker with respect to a root marker (the midpoint of the hip keypoints), normalized the 3D positions by the subject's height, sampled at 60 Hz, split the data into non-overlapping time-sequences of 0.5s, and added Gaussian noise (standard deviation: 0.018 m) to each time frame of the video keypoint positions based on a range of previously reported keypoint errors[13–15]. Finally, we standardized the data to have zero mean and unit standard deviation. We used the resulting time-sequences to train the networks.

We trained the LSTM networks in Python 3.7, using Keras 2.6, and one GPU (NVIDIA GeForce RTX 3090). The networks comprised LSTM layers followed by a dense layer with linear activation. We performed random searches of hyperparameters using the optimization framework Keras-Tuner[16] to select the number of LSTM layers, the number of units per LSTM layer, and the learning rate. We used the Adam gradient descent optimization algorithm[17] and the mean squared error as loss function. We selected hyperparameters that resulted in the lowest root mean squared error (RMSE) evaluated on the validation set and evaluated the performance of the networks on the test set using RMSE.

The random search of hyperparameters resulted in two (*body model*) and three (*arm model*) LSTM layers, 128 (*body model*) and 80 (*arm model*) units per LSTM layer, and learning rates of 7e-5 (*body model*) and 3e-5 (*arm model*). The RMSEs on the test set were 8.0 mm (*body model*) and 15.2 mm (*arm model*).

**Kinematic sensitivity analyses**

We used OpenCap to estimate anatomical marker locations, joint kinematics, ground reaction forces, and joint kinetics from videos. We evaluated the influence of the pose detection algorithm and the camera configuration on the estimated marker locations and joint kinematics.

We compared three pose detection algorithms/settings: HRNet[18–21] (person model: faster_rcnn_r50_fpn_coco, pose model: hrnet_w48_coco_wholebody_384x288_dark_plus), OpenPose[22] with default settings (later referred to as default OpenPose), and OpenPose with high accuracy settings (later referred to as high accuracy OpenPose). The default settings of OpenPose use a resolution of 208x368 pixels, whereas for the high accuracy settings we used a higher resolution (567x1008 pixels) than default and used OpenPose's scaling option that averages the results from processing the video at four different resolutions (selected resolution scaled by 1, 0.75, 0.5, and 0.25). The high accuracy settings result in more accurate marker position and kinematic estimates (see Table S1-S2) but require more GPU memory (>20 Gb) and more time for processing videos (about three times more than the default OpenPose settings). We also compared three camera configurations: five cameras (±70°, ±45°, and 0°), three cameras (±45° and 0°), and two cameras (±45°). The detailed results of these sensitivity analyses are presented in Table S1 (marker errors) and Table S2 (kinematic errors).

**Optimal control formulations**

OpenCap estimates kinetic measures using muscle-driven dynamic simulations that track 3D joint kinematics. These tracking simulations are formulated as optimal control problems.

We adjusted the musculoskeletal model and optimal control problem formulation, as compared to the generic formulation (see Methods: Design: Physics-based modeling and simulation), for the different activities to incorporate activity-based knowledge. First, we only included passive muscle forces for the walking simulations, as they were abnormally high for the other activities. This is expected as the musculoskeletal model was primarily validated based on walking, running, and pedaling data[23]. Second, we added reserve actuators (maximum of 30 Nm) to supplement the hip rotation muscle actuators for the squat and sit-to-stand simulations, as the model was otherwise too weak to track experimental kinematics with physiologically realistic muscle activations. This was also expected based on previous work[11]. Third, we added periodic constraints for the squat simulations. Squats are nearly periodic movements, and these constraints facilitate convergence of the optimal control problems. Note that these constraints are not necessary to obtain convergence. Finally, we added constraints for the squat and sit-to-stand simulations forcing the model to keep its heels in contact with the ground. We also added a similar term in the cost function for the sit-to-stand simulations to minimize the ratio between front-foot and rear-foot ground contact forces. These cost and constraint terms correspond to instructions given to participants during data collection to keep their feet flat on the ground. The terms also prevent the model from leaning forward to reduce muscle effort, which is modeled as squared muscle activations and minimized in the cost function.

**Calculation of knee loading measures**

We estimated the external knee adduction moment (KAM) and medial contact force (MCF) using the OpenSim API and Joint Reaction Force tool. To estimate the KAM, we replaced all muscles with ideal force and torque actuators at all degrees of freedom. At each time step, we posed the model, applied the ground reaction forces, and actuated the force and torque actuators to match the simulated or measured joint moments and pelvis residual forces and moments. We then performed a joint reaction analysis, and the KAM was considered the tibiofemoral reaction moment in the frontal plane of the tibia about the knee joint center. To estimate the MCF, we used the same procedure but actuated the model with the muscle forces from simulations instead of torque actuators. We then computed joint reaction forces and moments about the knee joint center. Assuming that contact forces in the medial and lateral compartments of the tibiofemoral joint balance the internal knee adduction moment, we computed MCF using Equation S1,

$$MCF = \frac{M_{\text{adduction}}}{d} + \frac{F_{\text{vertical}}}{2}, \tag{S1}$$

where $M_{\text{adduction}}$ is the internal tibiofemoral reaction moment in the frontal plane of the tibia about the knee joint center, $F_{\text{vertical}}$ is the tibiofemoral reaction force along the long axis of the tibia, and $d$ is the distance between the medial and lateral tibiofemoral contact points (assumed to be 4 cm[24]).

### Tables

Tables S1–S4 are included as independent files.

**Table S1: Errors in each marker position between OpenCap and motion capture.** The mean per marker error is shown for each marker, activity, camera combination, and pose detection algorithm.

**Table S2: Errors in kinematics between OpenCap and motion capture.** The mean absolute error (MAE) and root mean square error (RMSE) are shown for each degree of freedom, activity, camera combination, and pose detection algorithm.

**Table S3: Errors in ground reaction forces between OpenCap and force plates.** The mean absolute error (MAE), root mean square error (RMSE), and mean absolute error as a percentage of the range (MAPE) are shown for each activity using the two-camera HRNet setup.

**Table S4: Errors in joint moments between OpenCap and inverse dynamics using motion capture and force plates.** The mean absolute error (MAE), root mean square error (RMSE), and mean absolute error as a percentage of the range (MAPE) are shown for each activity and degree of freedom using the two-camera HRNet setup.

**Table S5: Statistical test information for walking case study.** Marker-based motion capture (Mocap) is compared to OpenCap using two cameras. All tests had nine degrees of freedom. For t tests, the test statistic is the t-score, the central tendency measure is the mean, the spread is the standard deviation, and the effect size is Cohen's d. For Wilcoxon signed rank test, the test statistic is W, the central tendency measure is the median, the spread is half of the interquartile range, the confidence interval is computed using bootstrap resampling (1000 samples), and the effect size is the common language effect size. Corrected *P*-values are reported after controlling for the false discovery rate.

| **Parameter** | **Test** | **Central tendency (spread)** | **95% Confidence interval** | **Test statistic** | **Corrected *P*-value** | **Effect size** |
|---|---|---|---|---|---|---|
| Change in peak knee adduction moment (%bodyweight*height) | | | | | | |
| OpenCap | t test | -0.89 (0.51) | (-1.27, -0.51) | -5.28 | .001 | 1.74 |
| Mocap | t test | -1.30 (0.54) | (-1.70, -0.89) | -7.27 | <.001 | 2.47 |
| Change in peak medial contact force (%bodyweight) | | | | | | |
| OpenCap | Wilcoxon | -49.6 (10.3) | (-65.5, -41.3) | 2 | .006 | 0.84 |
| Mocap | t test | -29.9 (30.4) | (-52.8, -7.0) | -2.95 | .016 | 1.14 |

**Table S6: Statistical test information for rising from a chair case study.** Marker-based motion capture (Mocap) is compared to OpenCap using two cameras. All tests had nine degrees of freedom. Corrected *P*-values are reported after controlling for the false discovery rate.

| **Parameter** | **Test** | **Mean (standard deviation)** | **95% Confidence interval** | **t-score** | **Corrected *P*-value** | **Effect size** |
|---|---|---|---|---|---|---|
| Change in knee extension moment (%bodyweight*height) | | | | | | |
| OpenCap | t test | -0.53 (0.58) | (-0.97, -0.09) | -2.71 | .024 | 0.88 |
| Mocap | t test | -1.04 (0.61) | (-1.51, -0.58) | -5.13 | .002 | 1.80 |
| Change in hip extension moment (%bodyweight*height) | | | | | | |
| OpenCap | t test | 0.75 (0.73) | (0.20, 1.30) | 3.07 | .020 | 1.30 |
| Mocap | t test | 1.00 (0.67) | (0.50, 1.50) | 4.50 | .002 | 1.70 |
| Change in ankle plantarflexion moment (%bodyweight*height) | | | | | | |
| OpenCap | t test | 1.05 (0.69) | (0.53, 1.57) | 4.59 | .004 | 1.80 |
| Mocap | t test | 1.23 (0.89) | (0.56, 1.91) | 4.15 | .003 | 2.13 |

### Figures

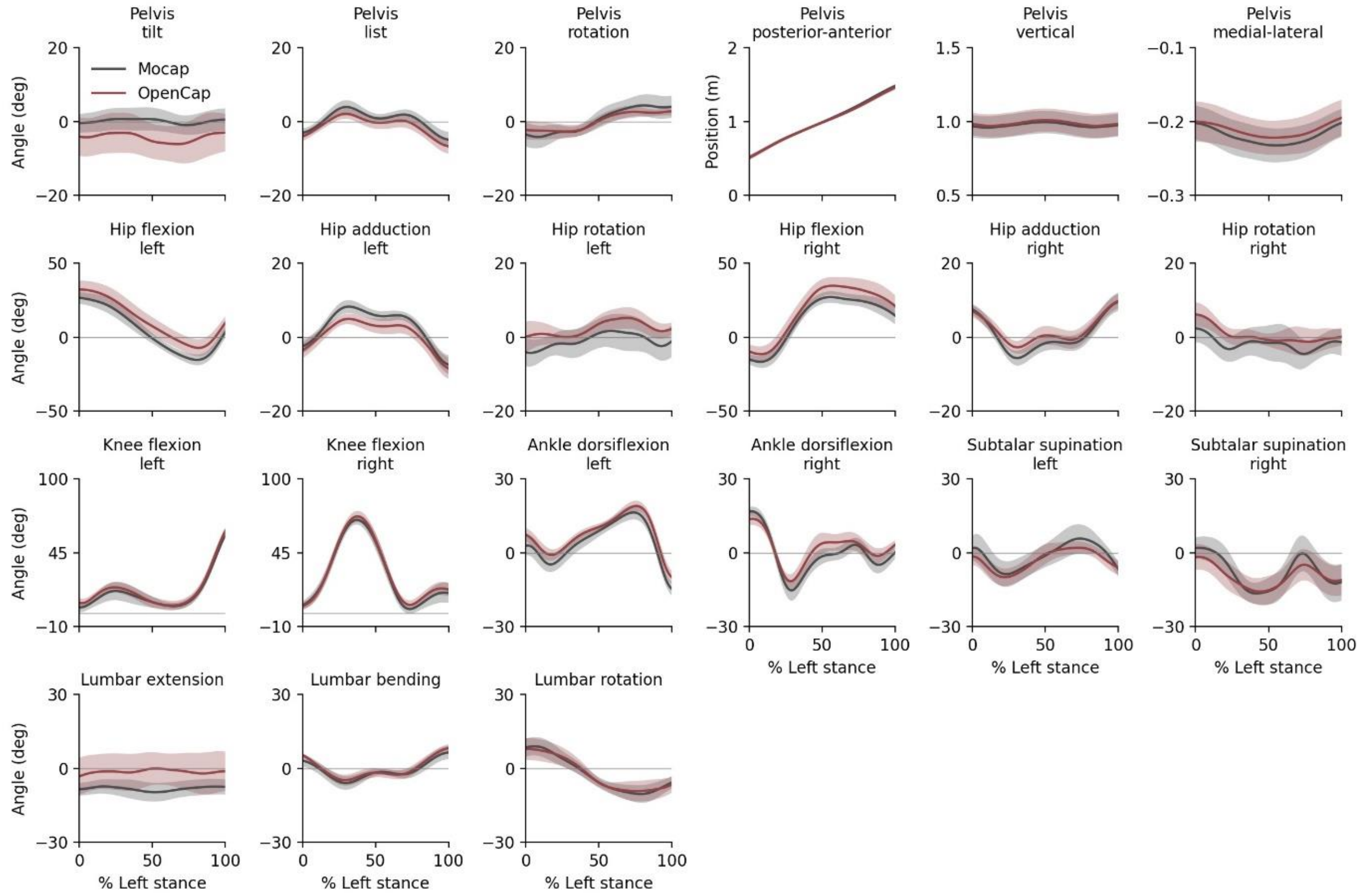


Figure S1: **Joint kinematics during natural walking**. The mean (line) and standard deviation (shading) across participants (n=10) of joint angles and positions estimated using OpenCap and based on marker-based motion capture (Mocap) are shown.

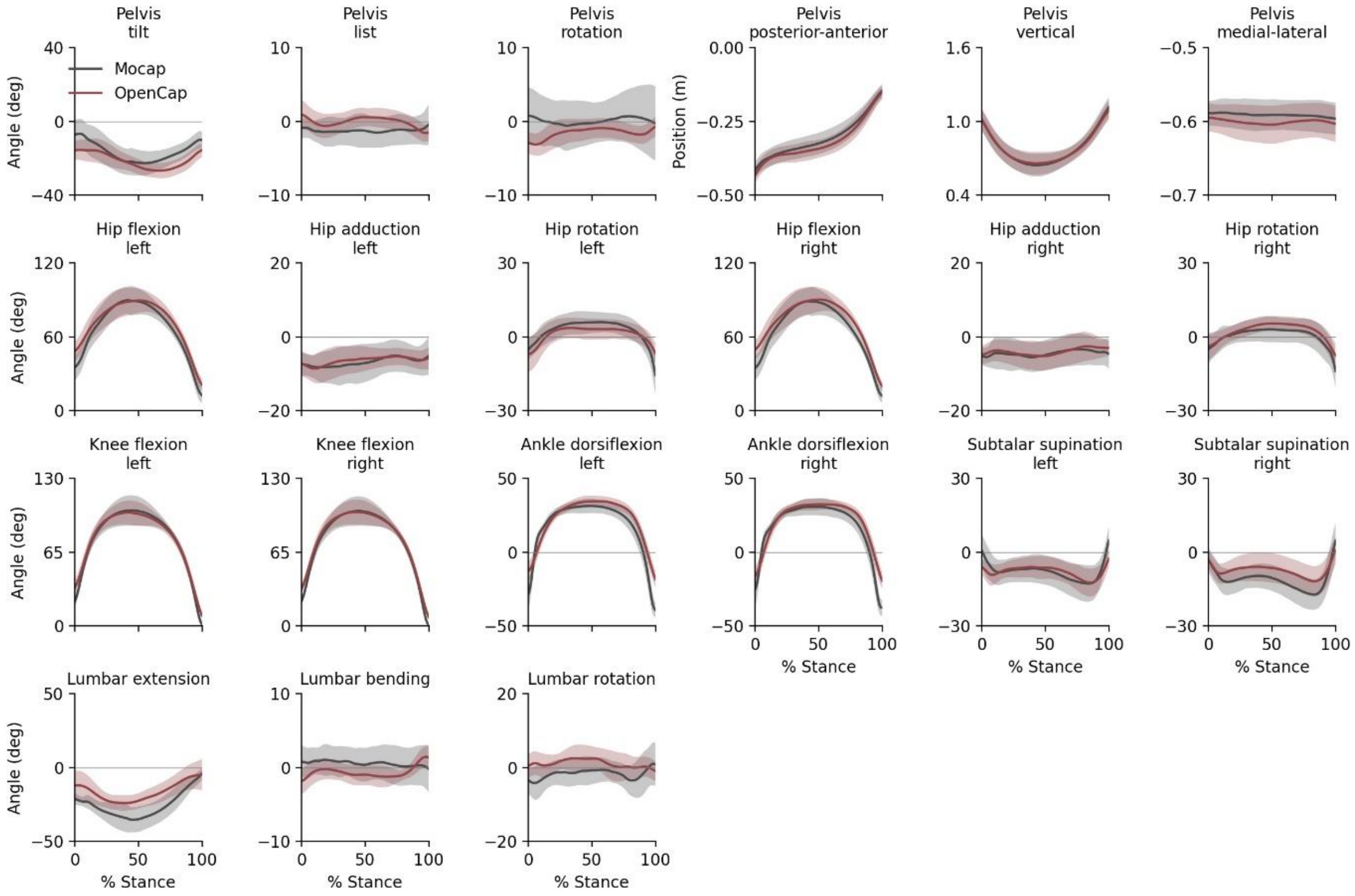


Figure S2: **Joint kinematics during natural drop jumps**. The mean (line) and standard deviation (shading) across participants (n=10) of joint angles and positions estimated using OpenCap and based on marker-based motion capture (Mocap) are shown.

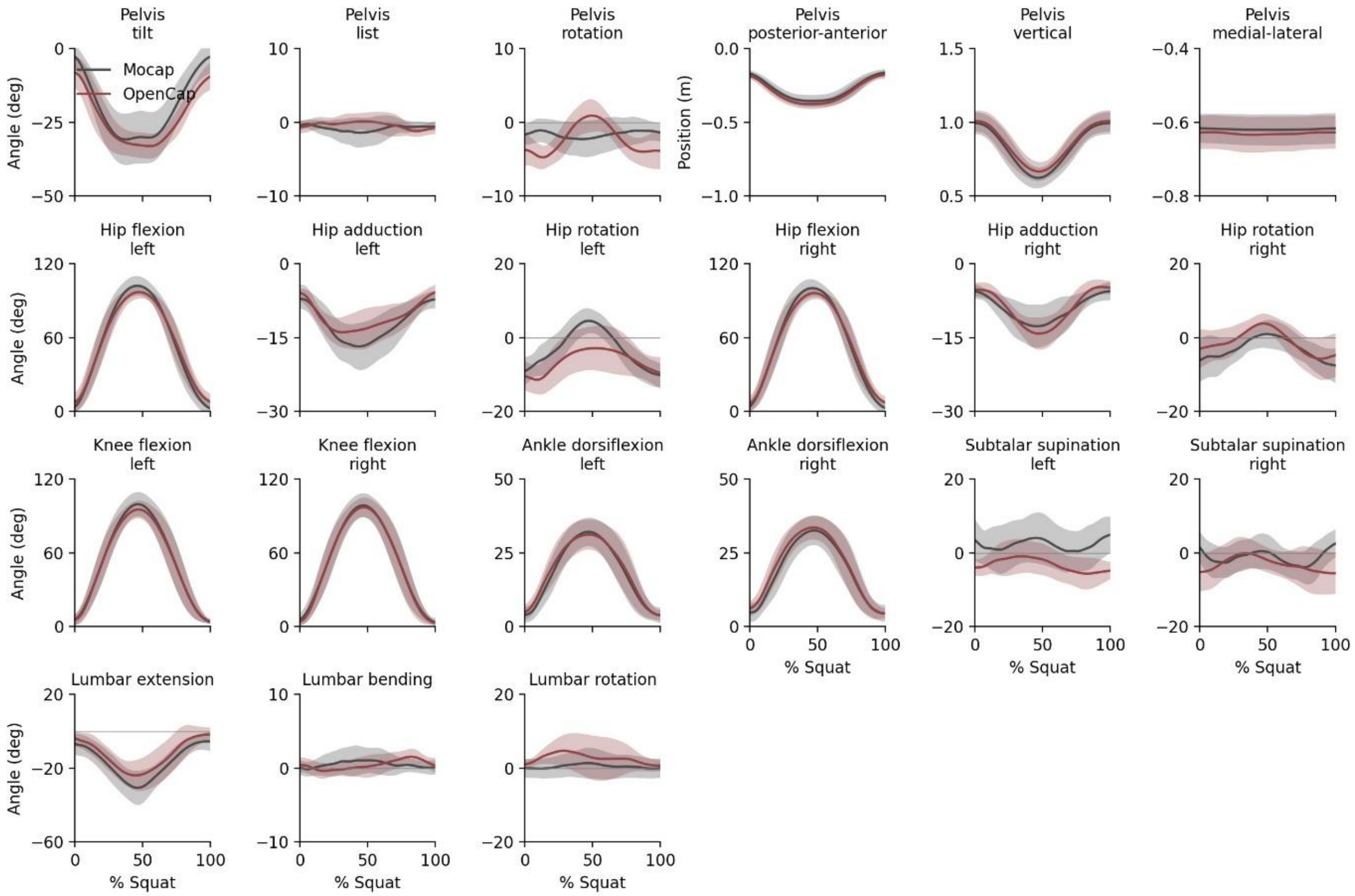


Figure S3: **Joint kinematics during natural squats**. The mean (line) and standard deviation (shading) across participants (n=10) of joint angles and positions estimated using OpenCap and based on marker-based motion capture (Mocap) are shown.

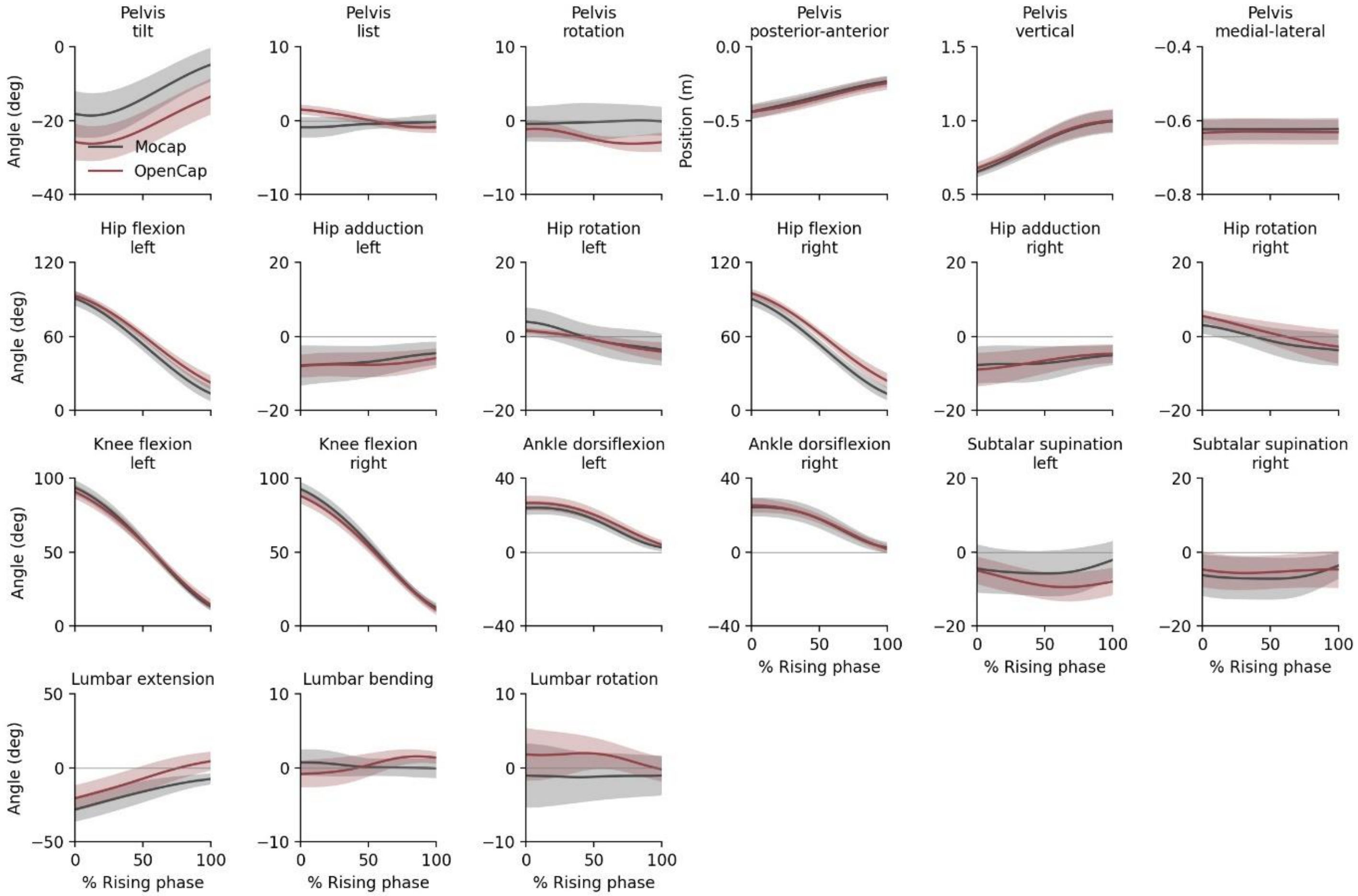


Figure S4: **Joint kinematics during natural sit-to-stands**. The mean (line) and standard deviation (shading) across participants (n=10) of joint angles and positions estimated using OpenCap and based on marker-based motion capture (Mocap) are shown.

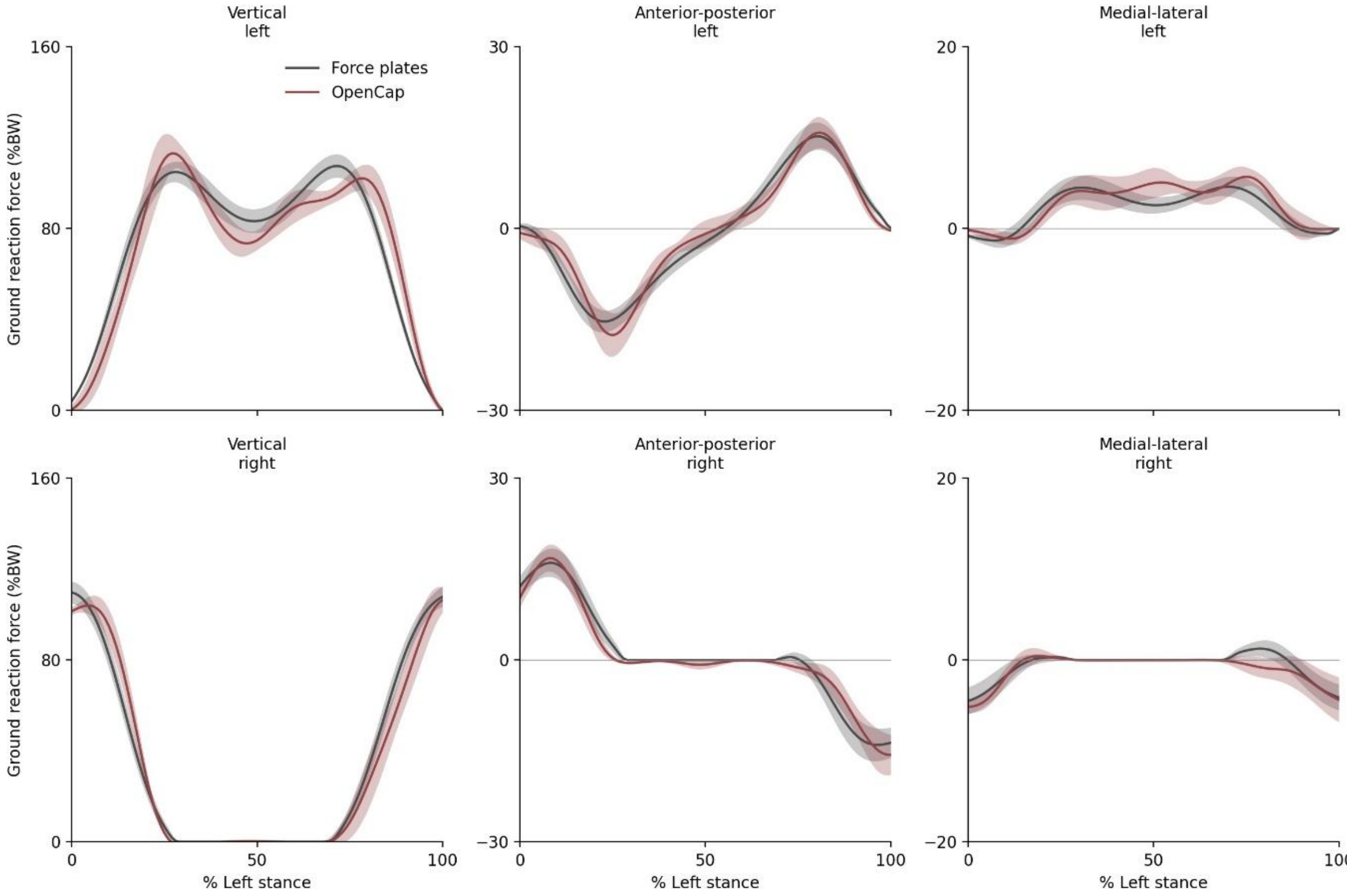


Figure S5: **Ground reaction forces during natural walking**. The mean (line) and standard deviation (shading) across participants (n=10) of ground reaction forces estimated using OpenCap and measured from force plates are shown. Forces are normalized to bodyweight (BW).

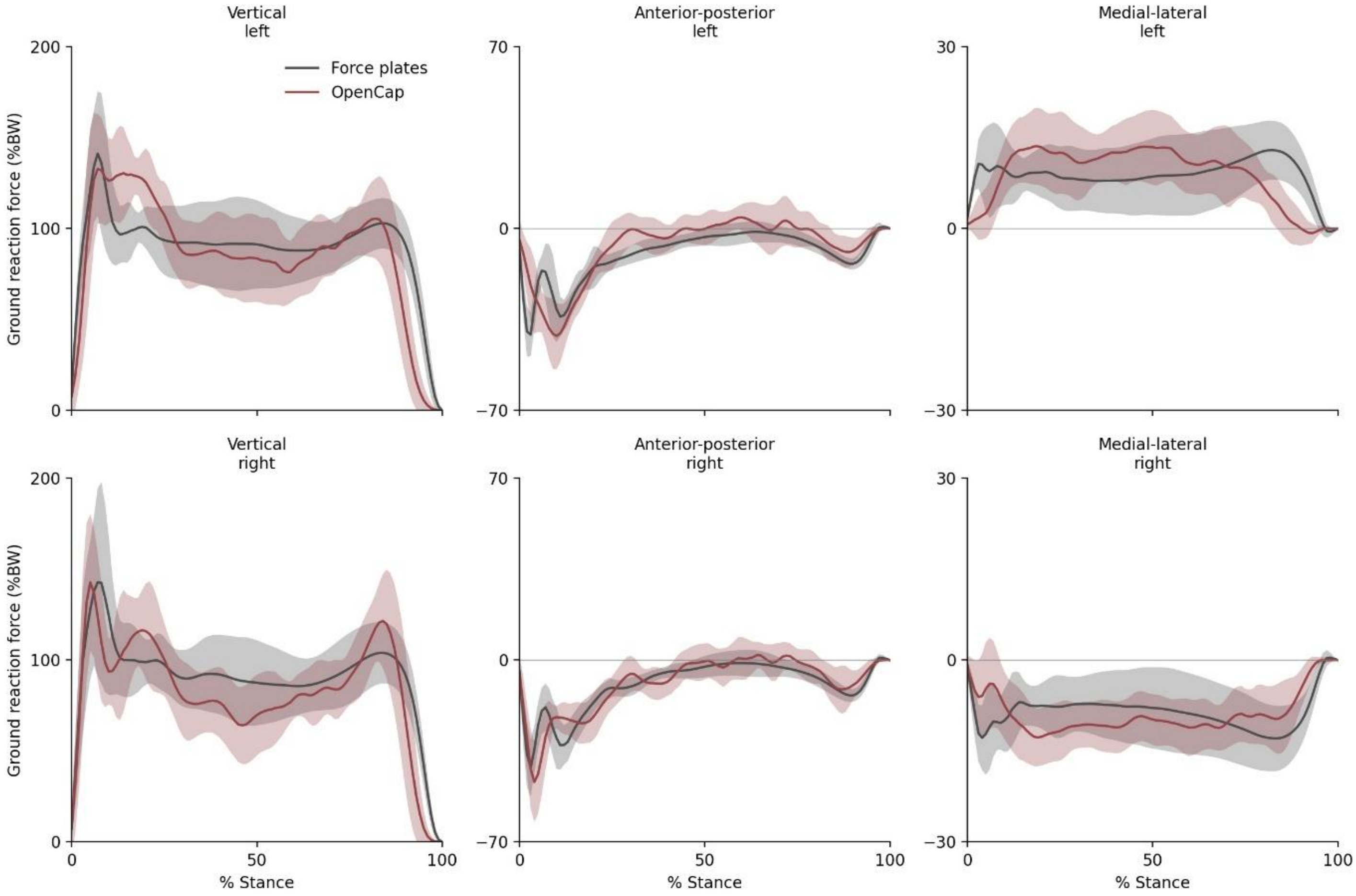


Figure S6: **Ground reaction forces during natural drop jumps**. The mean (line) and standard deviation (shading) across participants (n=10) of ground reaction forces estimated using OpenCap and measured from force plates are shown. Forces are normalized to bodyweight (BW).

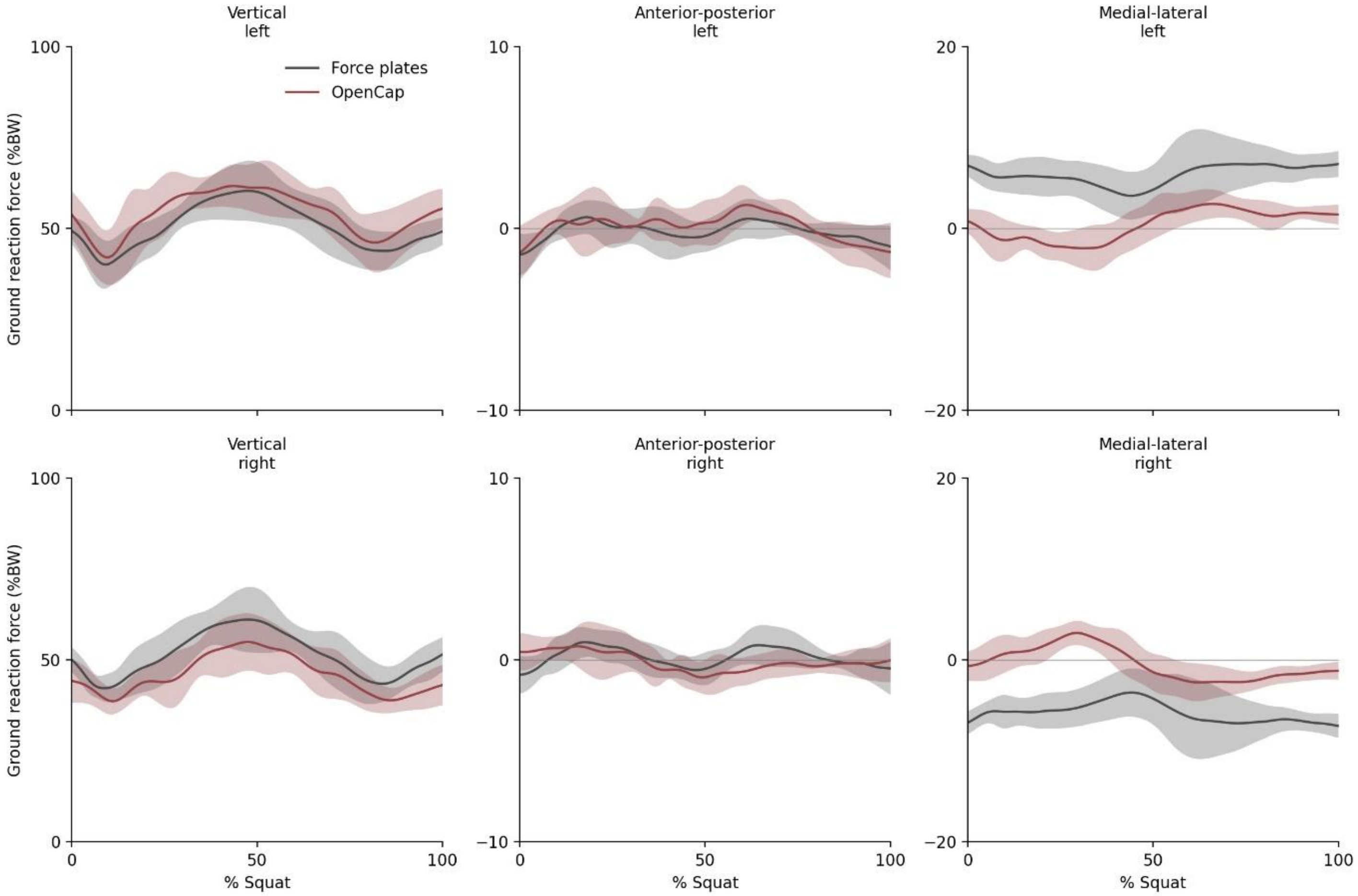


Figure S7: **Ground reaction forces during natural squats**. The mean (line) and standard deviation (shading) across participants (n=10) of ground reaction forces estimated using OpenCap and measured from force plates are shown. Forces are normalized to bodyweight (BW).

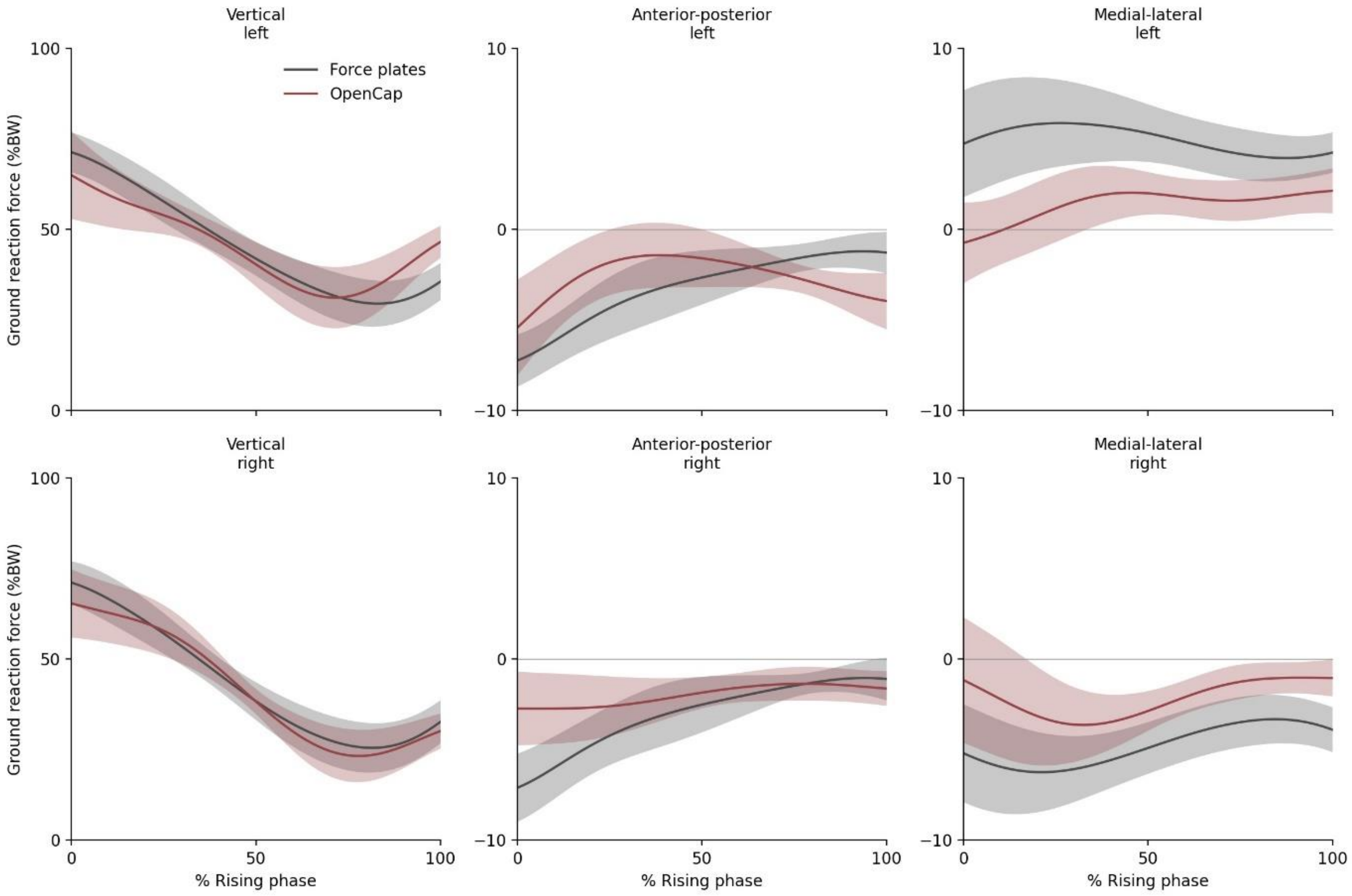


Figure S8: **Ground reaction forces during natural sit-to-stands**. The mean (line) and standard deviation (shading) across participants (n=10) of ground reaction forces estimated using OpenCap and measured from force plates are shown. Forces are normalized to bodyweight (BW).

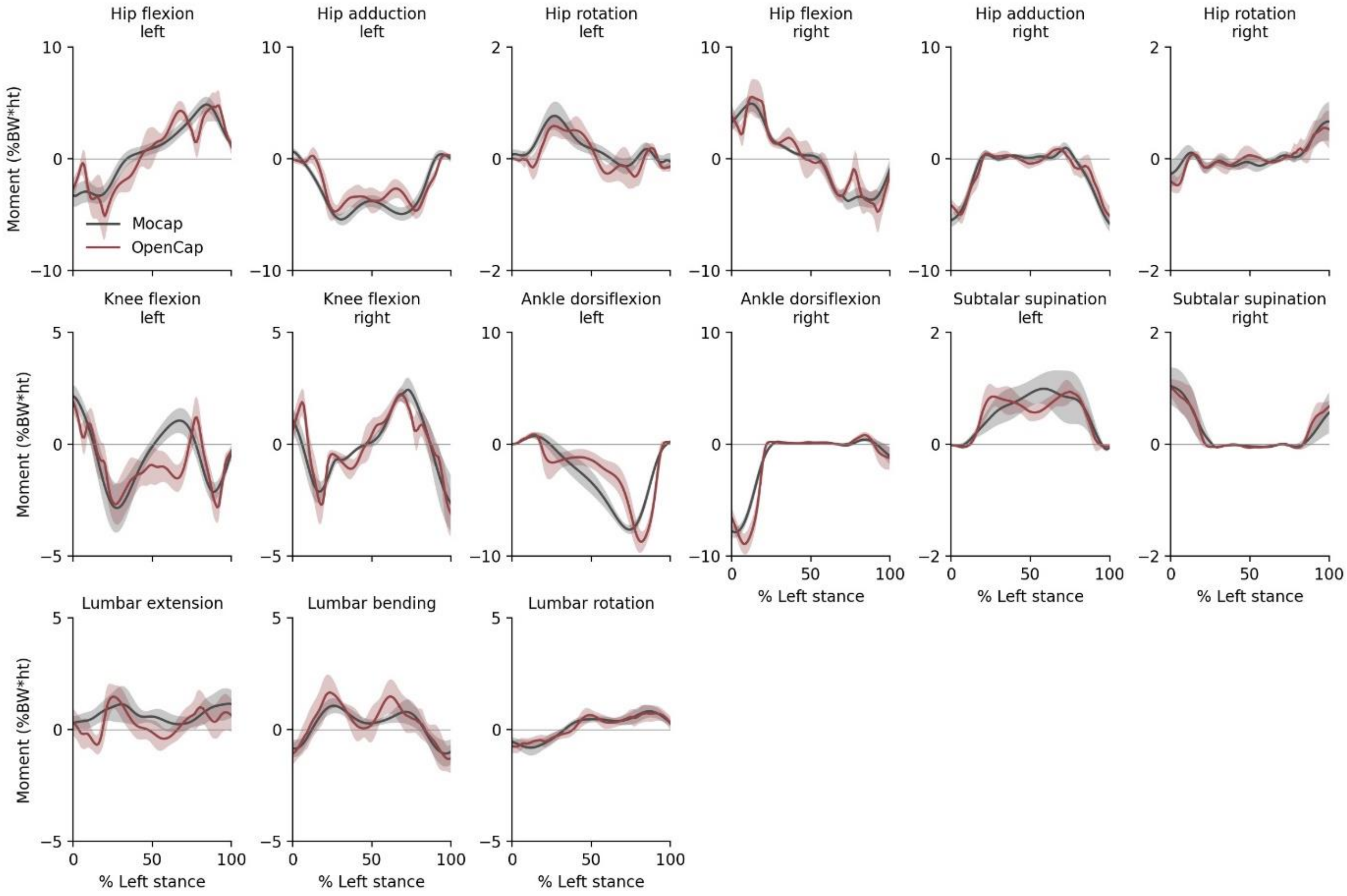


Figure S9: **Joint moments during natural walking**. The mean (line) and standard deviation (shading) across participants (n=10) of joint moments estimated using OpenCap and based on marker-based motion capture (Mocap) are shown. Moments are normalized to bodyweight (BW) and height (ht).

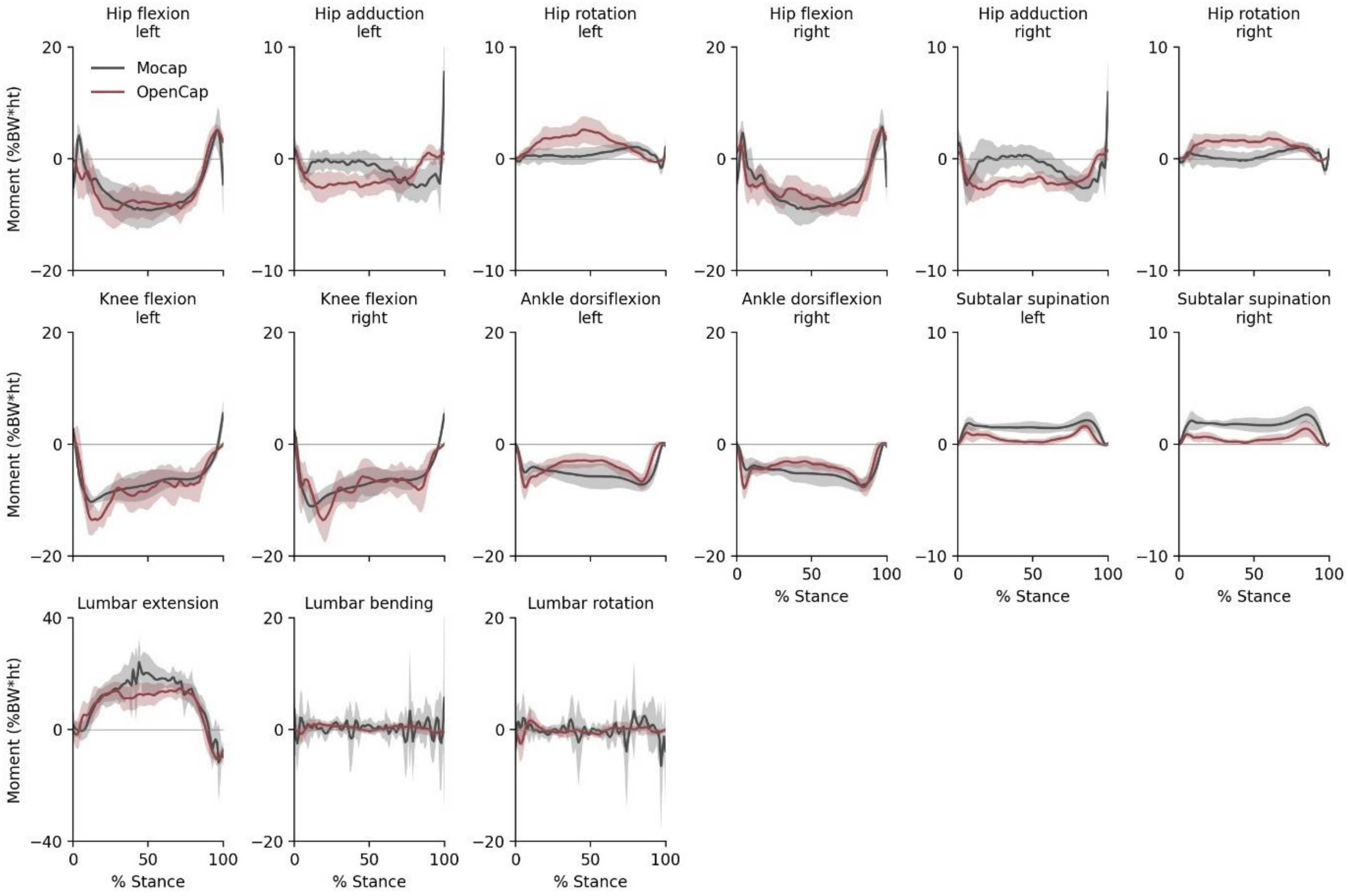


Figure S10: **Joint moments during natural drop jumps**. The mean (line) and standard deviation (shading) across participants (n=10) of joint moments estimated using OpenCap and based on marker-based motion capture (Mocap) are shown. Moments are normalized to bodyweight (BW) and height (ht).

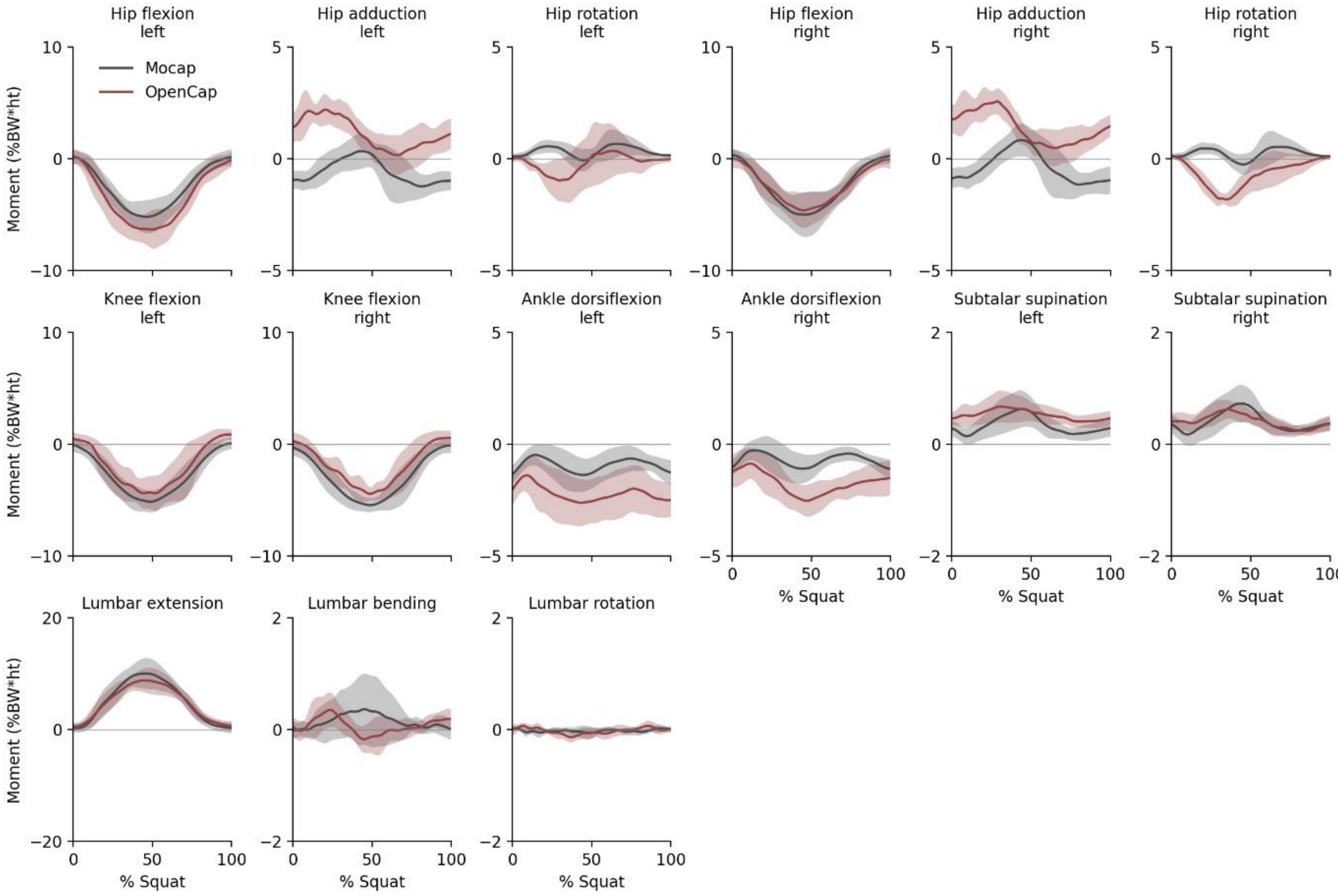


Figure S11: **Joint moments during natural squats**. The mean (line) and standard deviation (shading) across participants (n=10) of joint moments estimated using OpenCap and based on marker-based motion capture (Mocap) are shown. Moments are normalized to bodyweight (BW) and height (ht).

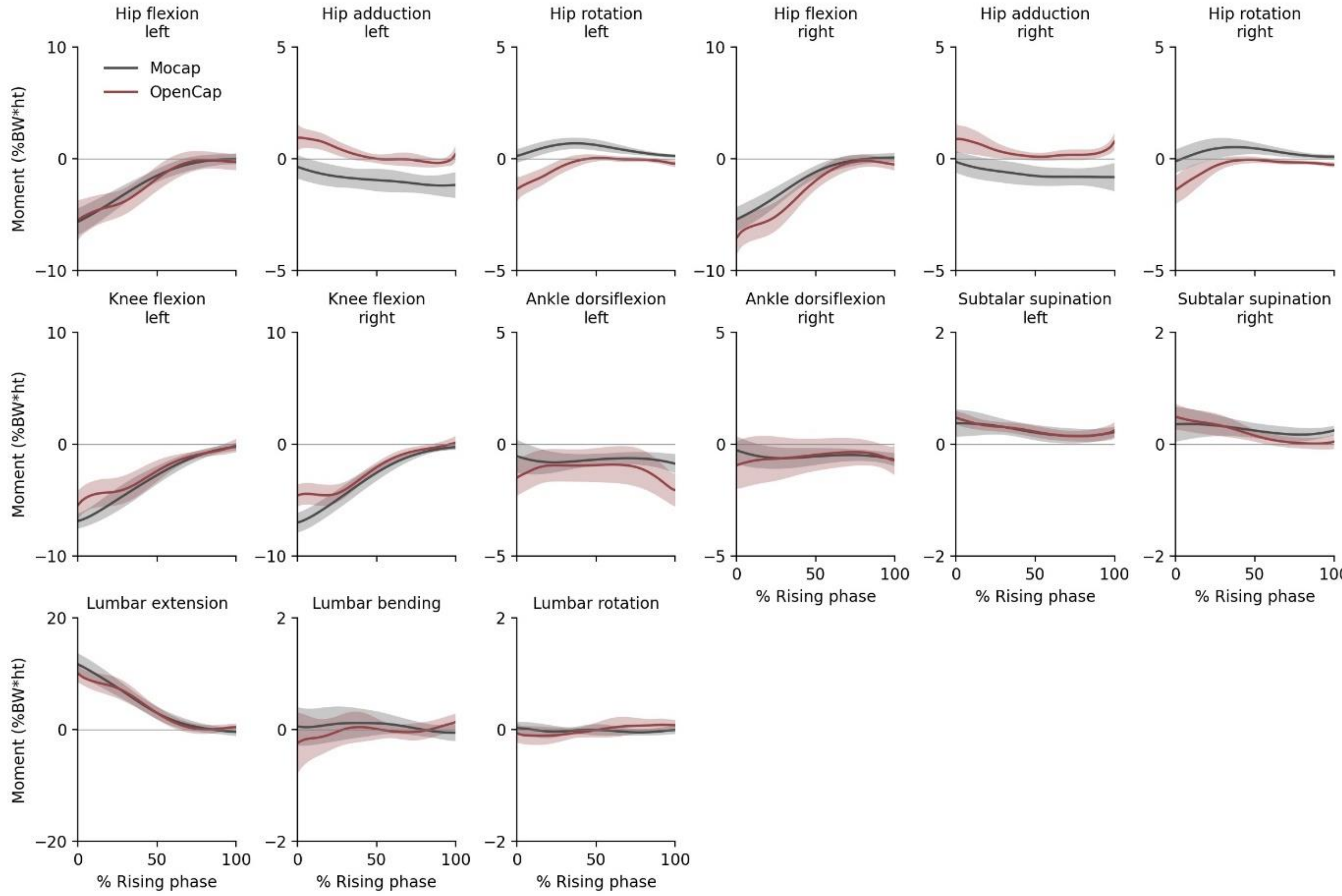


Figure S12: **Joint moments during natural sit-to-stands**. The mean (line) and standard deviation (shading) across participants (n=10) of joint moments estimated using OpenCap and based on marker-based motion capture (Mocap) are shown. Moments are normalized to bodyweight (BW) and height (ht).
